# Functional pathway redundancy in the metabolism of plant-derived phenolics by *Novosphingobium aromaticivorans*

**DOI:** 10.1101/2020.11.13.382549

**Authors:** Jose M. Perez, Wayne S. Kontur, Carson Gehl, Derek M. Gille, Yanjun Ma, Alyssa V. Niles, German Umana, Timothy J. Donohue, Daniel R. Noguera

## Abstract

Lignin is a plant heteropolymer composed of phenolic subunits. Because of its heterogeneity and recalcitrance, the development of efficient methods for its valorization still remains an open challenge. One approach to utilize lignin is its chemical deconstruction into mixtures of monomeric phenolic compounds followed by biological funneling into a single product. *Novosphingobium aromaticivorans* DSM12444 has been previously engineered to produce 2-pyrone-4,6-dicarboxylic acid (PDC) from depolymerized lignin by simultaneously metabolizing multiple aromatics through convergent routes involving the intermediates 3-methoxygallic acid (3-MGA) and protocatechuic acid (PCA). We investigated enzymes predicted to be responsible for *O*-demethylation and oxidative aromatic ring opening, two critical reactions involved in the metabolism of phenolics compounds by *N. aromaticivorans*. The results showed the involvement of DesA in *O*-demethylation of syringic and vanillic acids, LigM in *O-*demethylation of vanillic acid and 3-MGA, and a new *O-*demethylase, DmtS, in the conversion of 3-MGA into gallic acid (GA). In addition, we found that LigAB was the main aromatic ring opening dioxygenase involved in 3-MGA, PCA, and GA metabolism, and that a previously uncharacterized dioxygenase, LigAB2, had high activity with GA. Our results indicate a metabolic route not previously identified in *N. aromaticivorans* that involves *O*-demethylation of 3-MGA to GA. We predict this pathway channels ∼15% of the carbon flow from syringic acid, with the rest following ring opening of 3-MGA. The new knowledge obtained in this study allowed for the creation of an improved engineered strain for the funneling of aromatic compounds that exhibits stoichiometric conversion of syringic acid into PDC.

**IMPORTANCE:** For lignocellulosic biorefineries to effectively contribute to reduction of fossil fuel use, they need to become efficient at producing chemicals from all major components of plant biomass. Making products from lignin will require engineering microorganisms to funnel multiple phenolic compounds to the chemicals of interest, and *N. aromaticivorans* is a promising chassis for this technology. The ability of *N. aromaticivorans* to efficiently and simultaneously degrade many phenolic compounds may be linked to having functionally redundant aromatic degradation pathways and enzymes with broad substrate specificity. A detailed knowledge of aromatic degradation pathways is thus essential to identify genetic engineering targets to maximize product yields. Furthermore, knowledge of enzyme substrate specificity is critical to redirect flow of carbon to desired pathways. This study described an uncharacterized pathway in *N. aromaticivorans* and the enzymes that participate in this pathway, allowing the engineering of an improved strain for production of PDC from lignin.

## INTRODUCTION

Lignocellulosic plant biomass, composed of cellulose, hemicellulose, and lignin, is the most abundant organic material on the planet with potential to support a sustainable economy based on renewable feedstocks (1). Numerous studies predict that the economic and environmental viability of lignocellulosic biomass utilization for fuel and chemical production will be increased by the utilization of as much of these polymers as possible, including the use of lignin for production of chemicals (2-4). We are interested in deciphering the bacterial metabolism of phenolic compounds to engineer bacterial hosts to convert biomass-derived lignin into chemicals.

Lignin is an amorphous heteropolymer containing mainly syringyl (S; two methoxy groups), guaiacyl (G; one methoxy group), and *p*-hydroxyphenyl (H; no methoxy groups) phenolic structures that differ in the number of methoxy groups attached to the aromatic ring (5). One approach to valorizing lignin that has gained significant attention is to first use chemical techniques to deconstruct plant biomass and generate mixtures containing a large fraction of lower molecular weight phenolic compounds that could then be transformed by engineered microbes to a single valuable product (6). This funneling of phenolic mixtures to single compounds has been demonstrated with engineered strains of *Pseudomonas putida* (6, 7), *Rhodococcus jostii* (8), and *Novosphingobium aromaticivorans* (9). In addition, other microbes, such as the yeast *Rhodosporidium toruloides* (10) and the photoheterotrophic bacterium *Rhodopseudomonas palustris* (11-13) have been extensively studied for their ability to transform the plant-derived phenolic compounds often present in deconstructed plant biomass.

Among the desirable features for a microbial strain to be used as a chassis for the development of microbial lignin valorization strategies are an ability to metabolize the majority of the biomass-derived phenolic compounds and to funnel them into native convergent metabolic pathways (14). We are studying the sphingomonad bacterium *N. aromaticivorans* as a platform microbe for lignin valorization because it efficiently and simultaneously utilizes a large variety of S, G, and H phenolics (9, 15) and because it and other sphingomonads have enzymes to cleave different inter-subunit bonds in the lignin polymer (16-20). These features, when combined with the genetic and genomic information on sphingomonads like *N. aromaticivorans* could support strategies to maximize the number and type of lignin depolymerization products (e.g., phenolic monomers, oligomers) that can be microbially transformed into a range of valuable chemicals.

Metabolic pathways for the degradation of phenolics in sphingomonads have been proposed for *Sphingobium sp*. SYK-6, based on experiments with mutant strains and purified enzymes (21-23), and for *N. aromaticivorans*, based on analysis of a genome-scale transposon library and a set of targeted deletion mutants (9, 15). These studies have revealed several commonalities in the phenolic metabolism pathways of both organisms (Figure 1), but there remain significant knowledge gaps that limit the engineering of strains with increased transformation of phenolics to desired products.

**Figure 1.**
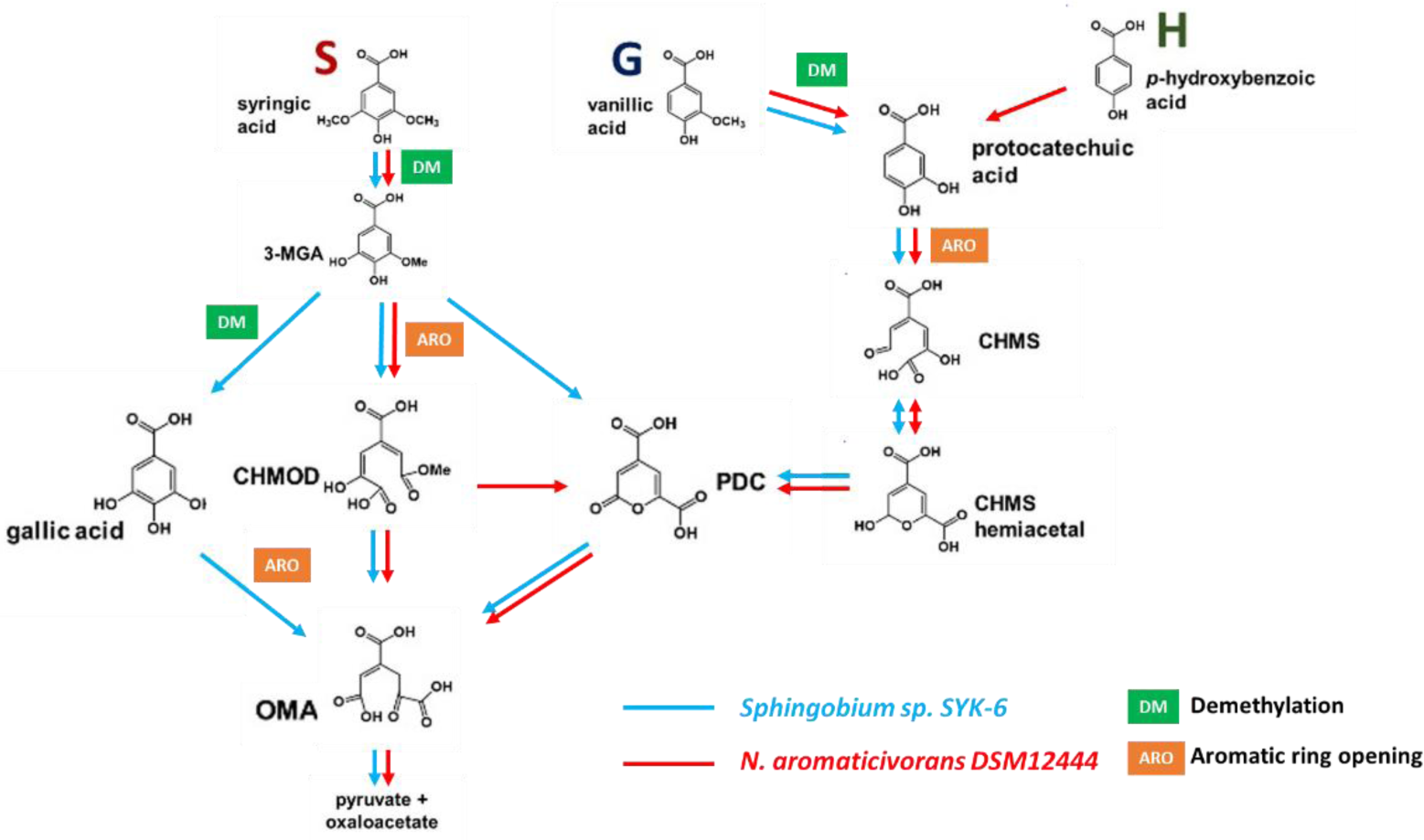
Pathways for the metabolism of S, G, and H phenolics that have been proposed for *Sphingobium sp*. SYK-6 (21-23) and *Novosphingobium aromaticivorans* (9, 15), showing the location of *O-*demethylation and aromatic ring opening steps.

To illustrate some of these knowledge gaps, syringic acid, which is an abundant biomass component that has been analyzed as a model S phenolic in both organisms, is demethylated to 3-methoxygallic acid (3-MGA), after which, multiple pathways have been proposed for 3-MGA transformation by *Sphingobium sp*. SYK-6 (21, 22). In one proposed pathway, the demethylation of 3-MGA produces gallic acid (GA), whose aromatic ring is cleaved by a dioxygenase to produce 4-oxalomesaconate (OMA) (24). Alternatively, it is proposed that the aromatic ring of 3-MGA is cleaved by a dioxygenase to produce 4-carboxy-2-hydroxy-6-methoxy-6-oxohexa-2,4-dienoate (CHMOD), which could be converted to OMA by an unknown enzyme (21, 22). Furthermore, *in vitro* experiments with a purified dioxygenase from *Sphingobium sp*. SYK-6 (LigAB) and 3-MGA as the substrate showed rapid production of 2-pyrone-4,6-dicarboxylic acid (PDC) in addition to the ring cleavage product CHMOD, a result that led to the hypothesis that LigAB catalyzes the transformation of 3-MGA to both CHMOD and PDC (22). In *N. aromaticivorans*, the only pathway thus far proposed for syringic acid metabolism (15) is via demethylation to 3-MGA, ring cleavage to CHMOD, and conversion to OMA. However, a *N. aromaticivorans* deletion mutant lacking the proposed enzymes for CHMOD conversion to OMA (DesC and DesD) resulted in accumulation of PDC, suggesting the possibility for cyclization of CHMOD to PDC that is independent of these enzymes (9).

There are also knowledge gaps in the bacterial metabolism of the other major G and H phenolic substituents of plant cell walls. In the case of the G phenolic vanillic acid, its metabolism by both *Sphingobium* sp. SYK-6 and *N. aromaticivorans* is proposed (15, 16) to entail demethylation to protocatechuic acid (PCA), ring cleavage to 4-carboxy-2-hydroxy-*cis,cis*-muconate-6-semialdehyde (CHMS), oxidation to PDC, and hydrolysis to OMA (Figure 1). Degradation of the H phenolic *p*-hydroxybenzoic acid has only been studied in *N. aromaticivorans* (9), with experimental evidence suggesting transformation to PCA as the initial step to enter the described pathway for G phenolics (Figure 1). However, it is unclear if the S and G/H branches of the phenolic degradation pathways in sphingomonads have common or pathway-specific *O-*demethylation and aromatic ring cleavage enzymes (Figure 1). Indeed, it has been proposed that some of these demethylase enzymes have broad substrate specificity and are active in multiple branches. For example, one dioxygenase (LigAB) in *N. aromaticivorans* has been proposed to be active in the ring cleavage of 3-MGA and PCA (15), and in *Sphingobium sp*. SYK-6, an *O-*demethylase (LigM) has been proposed to be active on both 3-MGA and vanillic acid (25).

To address knowledge gaps on the reactions and enzymes involved in S, G, and H phenolic metabolism by *N. aromaticivorans*, we analyzed putative *O-*demethylases and aromatic ring opening dioxygenases that are predicted to be involved in metabolism of these plant-derived aromatics. We present results of experiments with purified enzymes and with deletion mutants that provide evidence for functionally redundant pathways and for enzymes with broad substrate specificity in this organism. Our studies led us to identify an aromatic ring-opening dioxygenase (LigAB2) and an *O-*demethylase (DmtS), which have not been previously shown to have these activities in sphingomonads. In addition, the newly acquired knowledge on enzyme redundancy and substrate specificity allowed us to engineer a second-generation *N. aromaticivorans* DSM 12444 strain with improved yields of PDC from plant-derived phenolics.

## RESULTS

### Identification of putative aromatic *O-*demethylases in *N. aromaticivorans*

We evaluated gene products with significant amino acid sequence identity to *O-*demethylases, encoded by Saro_2861 (*ligM*) and Saro_2404 (*desA*), that have been proposed to be involved in vanillic and syringic acid metabolism, respectively (15). These two proteins share ∼78% and ∼71% amino acid sequence identity with the *Sphingobium sp*. SYK-6 *O-*demethylases SLG_12740 (*ligM*) (25) and SLG_25000 (*desA*) (26). In addition, since *O-*demethylation of vanillic acid in *Pseudomonas* (27, 28) is catalyzed by VanAB, we analyzed the *N. aromaticivorans* gene Saro_1872 (hereafter called *dmtS*), which encodes a protein with the closest amino acid sequence identity to the VanA subunit of this enzyme (e.g., ∼27% amino acid identity with VanA of *Pseudomonas sp*. strain HR199 (29)).

### Effect of deleting putative O-demethylase genes on N. aromaticivorans growth

To begin evaluating the involvement of *ligM, desA*, and *dmtS* in the degradation of S and G phenolics in *N. aromaticivorans* we generated mutants (Table 1) containing combinations of deletions of these 3 genes in a parent strain (12444*Δ1879*) and in a strain (12444PDC) in which deletions in PDC- and CHMOD-degradation genes led to accumulation of PDC from S, G, and H aromatics (9).

**Table 1.** List of *N. aromaticivorans* parent and mutant strains with deletions of putative *O-*demethylases used in this study.

| Strain | Background used for strain construction | Relevant Characteristics |
| --- | --- | --- |
| Parent strain (12444Δ1879)* | DSM12444 | 12444Δ1879 |
| PDC-producing strain (12444PDC)* | 12444Δ1879 | 12444Δ1879 ΔSaro_2819 ΔSaro_2864 ΔSaro_2865 |
| 12444Δ*ligM* | 12444Δ1879 | 12444Δ1879 ΔSaro_2861 |
| 12444Δ*desA* | 12444Δ1879 | 12444Δ1879 ΔSaro_2404 |
| 12444Δ*dmrS* | 12444Δ1879 | 12444Δ1879 ΔSaro_1872 |
| 12444Δ*ligMΔdesA* | 12444Δ*ligM* | 12444Δ1879 ΔSaro_2861 ΔSaro_2404 |
| 12444Δ*ligMΔdmrS* | 12444Δ*ligM* | 12444Δ1879 ΔSaro_2861 ΔSaro_1872 |
| 12444Δ*desAΔdmrS* | 12444Δ*dmrS* | 12444Δ1879 ΔSaro_2404 ΔSaro_1872 |
| 12444Δ*ligMΔdesAΔdmrS* | 12444Δ*ligMΔdmrS* | 12444Δ1879 ΔSaro_2861 ΔSaro_2404 ΔSaro_1872 |
| 12444PDCΔ*ligM* | 12444PDC | 12444PDC ΔSaro_2861 |
| 12444PDCΔ*desA* | 12444PDC | 12444PDC ΔSaro_2404 |
| 12444PDCΔ*dmrS* | 12444PDC | 12444PDC ΔSaro_1872 |
| 12444PDCΔ*ligMΔdesA* | 12444PDCΔ*ligM* | 12444PDC ΔSaro_2861 ΔSaro_2404 |
| 12444PDCΔ*ligMΔdmrS* | 12444PDCΔ*ligM* | 12444PDC ΔSaro_2861 ΔSaro_1872 |
| 12444PDCΔ*desAΔdmrS* | 12444PDCΔ*dmrS* | 12444PDC ΔSaro_2404 ΔSaro_1872 |
| 12444PDCΔ*ligMΔdesAΔdmrS* | 12444PDCΔ*ligMΔdmrS* | 12444PDC ΔSaro_2861 ΔSaro_2404 ΔSaro_1872 |

\*Construction of the parent and PDC-producing strains is described in Perez et al. (2019) (9). The parent strain has a deletion of *Saro\_1879 (sacB)* to render it susceptible to genetic manipulation using *sacB* as a countersensitive marker (9).

Figure 2 shows growth curves for the parent strain and for its corresponding set of mutant strains. When cultured in the presence of only syringic acid, the parent strain and the mutant strains 12444*ΔligM*, 12444*ΔdmtS*, and 12444*ΔligMΔdmtS* had similar growth patterns, whereas all mutant strains lacking *desA* were unable to grow (Figure 2A). This suggests that *desA* is essential for *N. aromaticivorans* growth on syringic acid, whereas *ligM* and *dmtS* are not.

**Figure 2.**
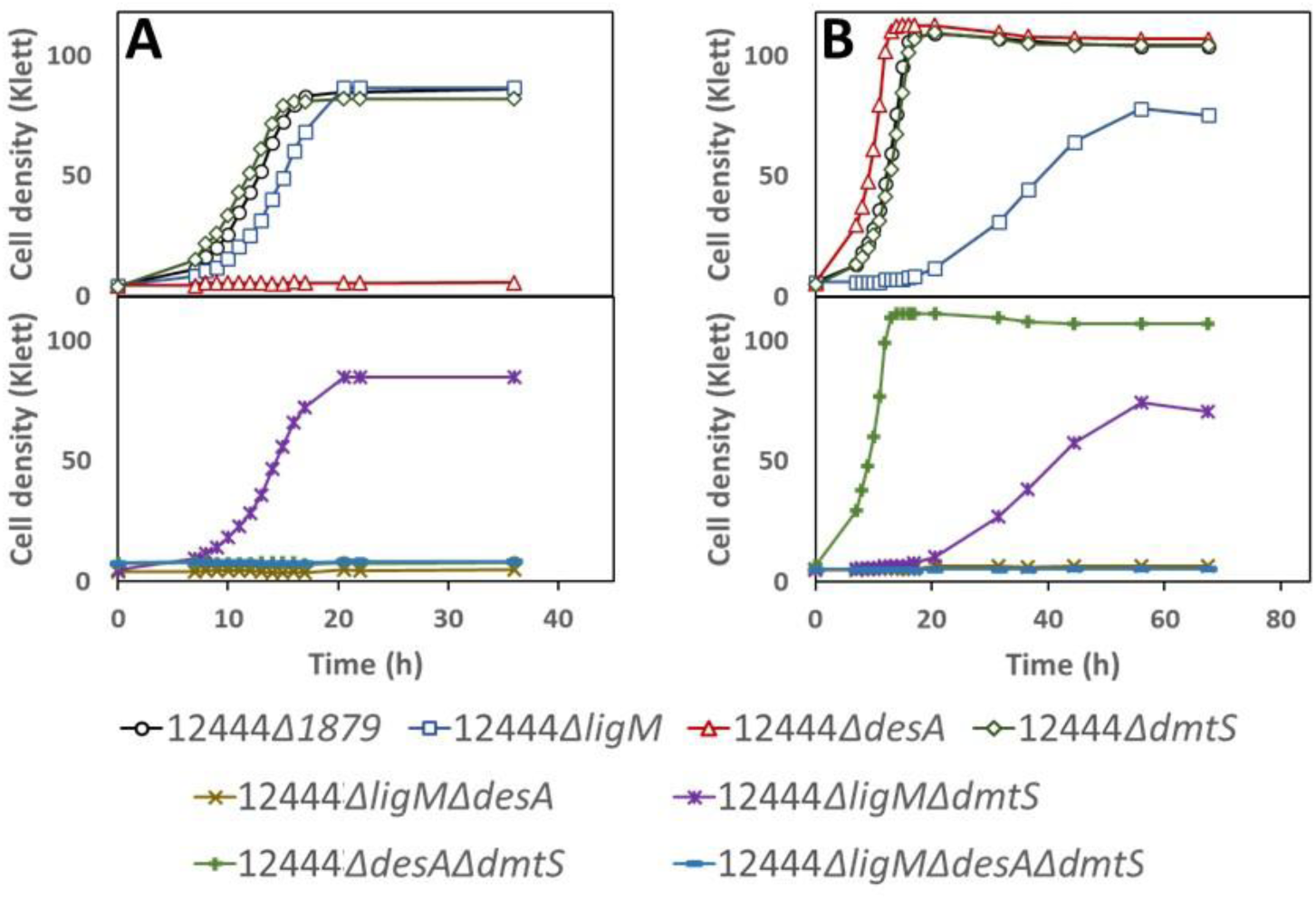
Growth of indicated strains of *N. aromaticivorans* in minimal media supplemented with syringic acid (panel A) or vanillic acid (panel B). Top panels show data for parent strain and single deletion mutants, whereas bottom panels show growth curves for strains with multiple gene deletions.

When cultured in the presence of vanillic acid as the sole carbon source, three distinct growth patterns were observed for this set of mutants (Figure 2B). First, all mutant strains with an intact *ligM* (12444*ΔdesA*, 12444*ΔdmtS*, and 12444*ΔdesAΔdmtS*) exhibited a similar growth pattern as the parent strain, suggesting that the DesA and DmtS enzymes are not essential for *N. aromaticivorans* growth on vanillic acid. Second, strains 12444*ΔligM* and 12444*ΔligMΔdmtS* showed lower growth rates and a lower final cell density compared to the parent strain, and third, strains simultaneously lacking *desA* and *ligM*, 12444*ΔdesAΔligM* and 12444*ΔdesAΔligMΔdmtS* were unable to grow in the presence of vanillic acid.

Because the growth defects caused by gene deletions in the parent strain do not necessarily reveal which specific intracellular reactions are being affected, and can obscure the roles of genes with redundant functions, we also used the PDC-producing strain as a background to construct mutants lacking the same combinations of putative *O-*demethylase genes (Table 1). With this second set of mutants, which require a non-aromatic carbon source for growth, we used PDC production as a proxy to elucidate the metabolic pathways affected by the gene deletions.

In the presence of glucose and syringic acid as an aromatic carbon source, the original PDC-producing strain (12444PDC) completely removed the syringic acid from the medium and produced PDC with a yield of ∼85% (Figure 3A; Table 2). Strains with deletions of *desA* plus at least one more putative *O-* demethylase (12444PDC*ΔdesAΔligM*, 12444PDC*ΔdesAΔdmtS*, and 12444PDC*ΔdesAΔligMΔdmtS*) were not able to degrade syringic acid (Figure 3E, 3F, 3H), whereas the strain with the single *desA* deletion consumed syringic acid and produced PDC at a ∼88% yield (Figure 3B; Table 2). As three out of the four mutants containing a deletion of *desA* were unable to consume syringic acid, these results support a role for DesA in syringic acid metabolism by *N. aromaticivorans*, which is proposed to be its demethylation to 3-MGA (Figure 1). However, PDC production by single mutants lacking only the *desA* gene suggests that, when *N. aromaticivorans* is not depending on aromatics for growth, LigM and DmtS may substitute for DesA activity in the metabolism of syringic acid.

**Figure 3.**
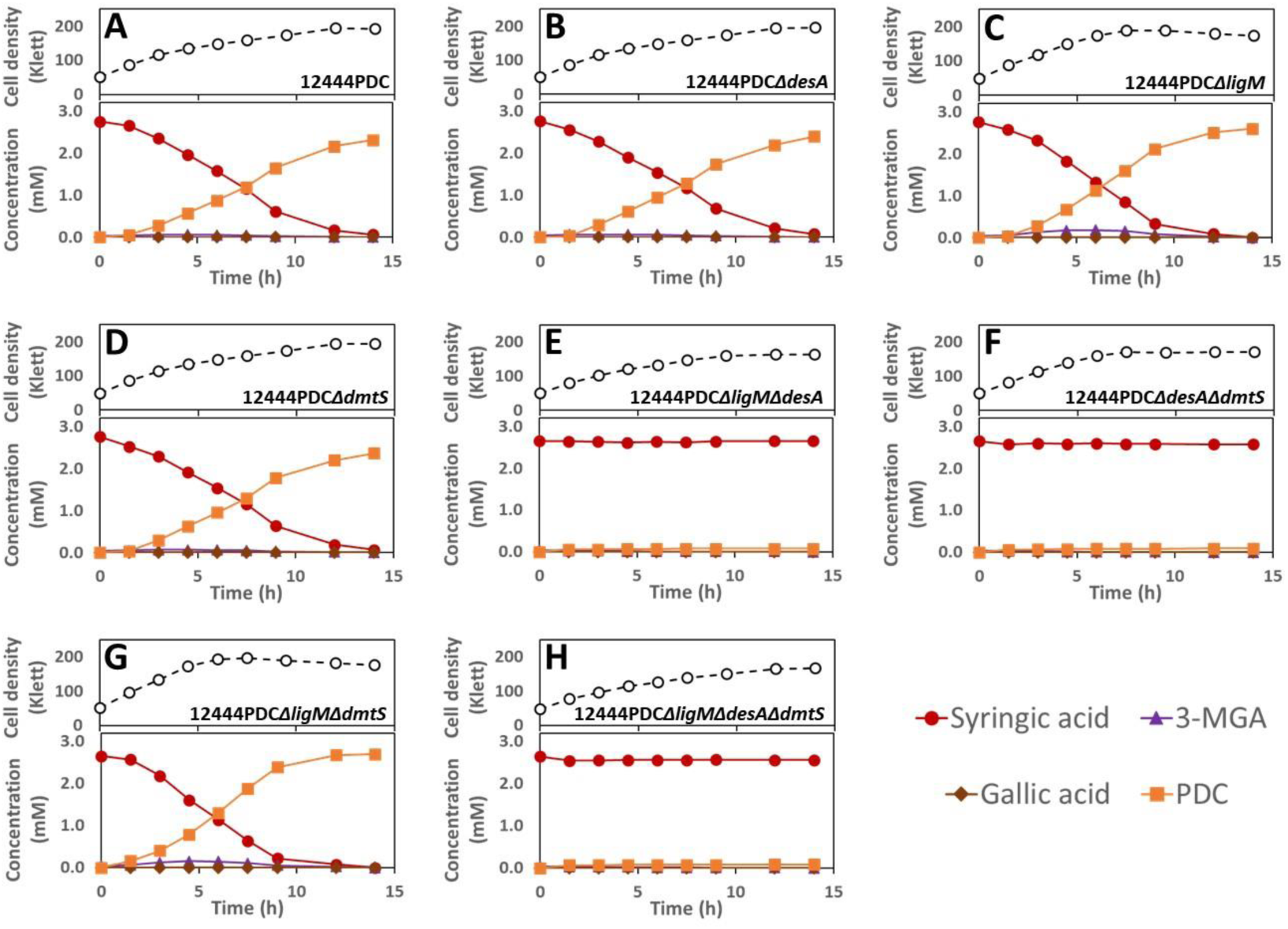
Cell densities and extracellular compound concentrations for the indicated *N. aromaticivorans* strains grown in minimal media containing glucose and syringic acid.

**Table 2.**
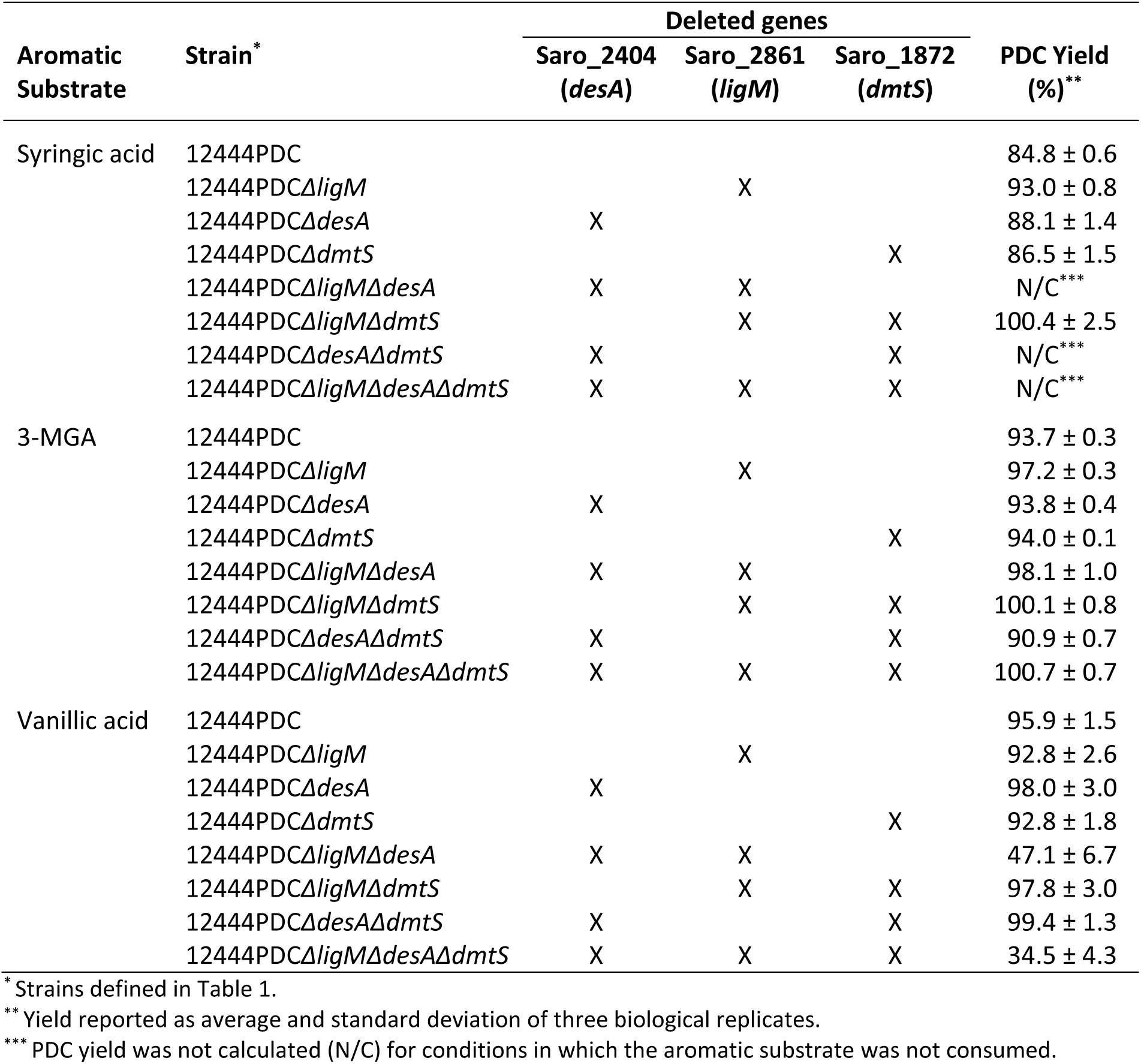
PDC yields from *N. aromaticivorans* strains in the presence of glucose plus the indicated aromatic substrate.

Furthermore, all the mutant strains with intact *desA* that were constructued in the PDC-producing background (12444PDC*ΔligM*, 12444PDC*ΔdmtS*, and 12444PDC*ΔligMΔdmtS*) consumed syringic acid and accumulated PDC. Notably, the PDC yield from syringic acid by strain 12444PDC*ΔligMΔdmtS* was stoichiometric and reproducibly higher than the yield of this product obtained in the other strains (Table 2). This suggests that the simultaneous deletion of both *ligM* and *dmtS* blocks another previously unknown pathway for syringic acid metabolism that normally detracts from PDC production in the 12444PDC strain. Since the PDC yield by the 12444PDC strain was ∼85%, we interpret these results as indicating that only a small fraction of the syringic acid is metabolized via this previously unidentified pathway.

We predict that this alternative pathway for syringic acid metabolism involves *O-*demethylation (by LigM and/or DmtS) of 3-MGA to GA, since that pathway has been proposed to occur in *Sphingobium sp*. SYK-6 (Figure 1). An observed small and transient accumulation of 3-MGA in the culture media when 12444PDC*ΔligM* and 12444PDC*ΔligMΔdmtS* (Figure 3C, 3G) are grown in the presence of syringic acid supports this hypothesis. To further tests this hypothesis, we grew the 12444PDC strain and its corresponding set of putative *O-*demethylase deletion mutants in the presence of glucose and 3-MGA (Figure 4). We posited that using 3-MGA as the aromatic substrate would provide a growth substrate that only requires one demethylation reaction compared to growth with syringic acid that is predicted to require the removal of two methyl groups. In this set of experiments, strain 12444PDC was able to completely remove 3-MGA from the medium, converting ∼94% of it into PDC. Moreover, in contrast with the experiments using syringic acid (Figure 3), each of the putative *O-*demethylase mutant derivatives of 12444PDC were able to completely remove 3-MGA from the media (Figure 4). This result indicates that none of these putative *O-*demethylases is essential for conversion of 3-MGA to PDC. It also suggests that 3-MGA is primarily metabolized via ring cleavage in *N. aromaticivorans*, a process that does not require *O-*demethylation (Figure 1). In addition, all strains with deletions of *ligM* (12444PDC*ΔligM*, 12444PDC*ΔdesAΔligM*, 12444PDC*ΔligMΔdmtS*, and 12444PDC*ΔdesAΔligMΔdmtS*) reproducibly produced more PDC than strain 12444PDC (Table 2), a result that suggests the involvement of LigM in the proposed alternative pathway of syringic acid metabolism. Notably, stoichiometric conversion of 3-MGA to PDC was achieved in the mutants with simultaneous deletion of *ligM* and *dmtS* (12444PDC*ΔligMΔdmtS*, and 12444PDC*ΔdesAΔligMΔdmtS*), which, in agreement with the experiments using syringic acid, suggests that the loss of LigM and DmtS blocks the alternative pathway for syringic acid metabolism in this bacterium.

**Figure 4.**
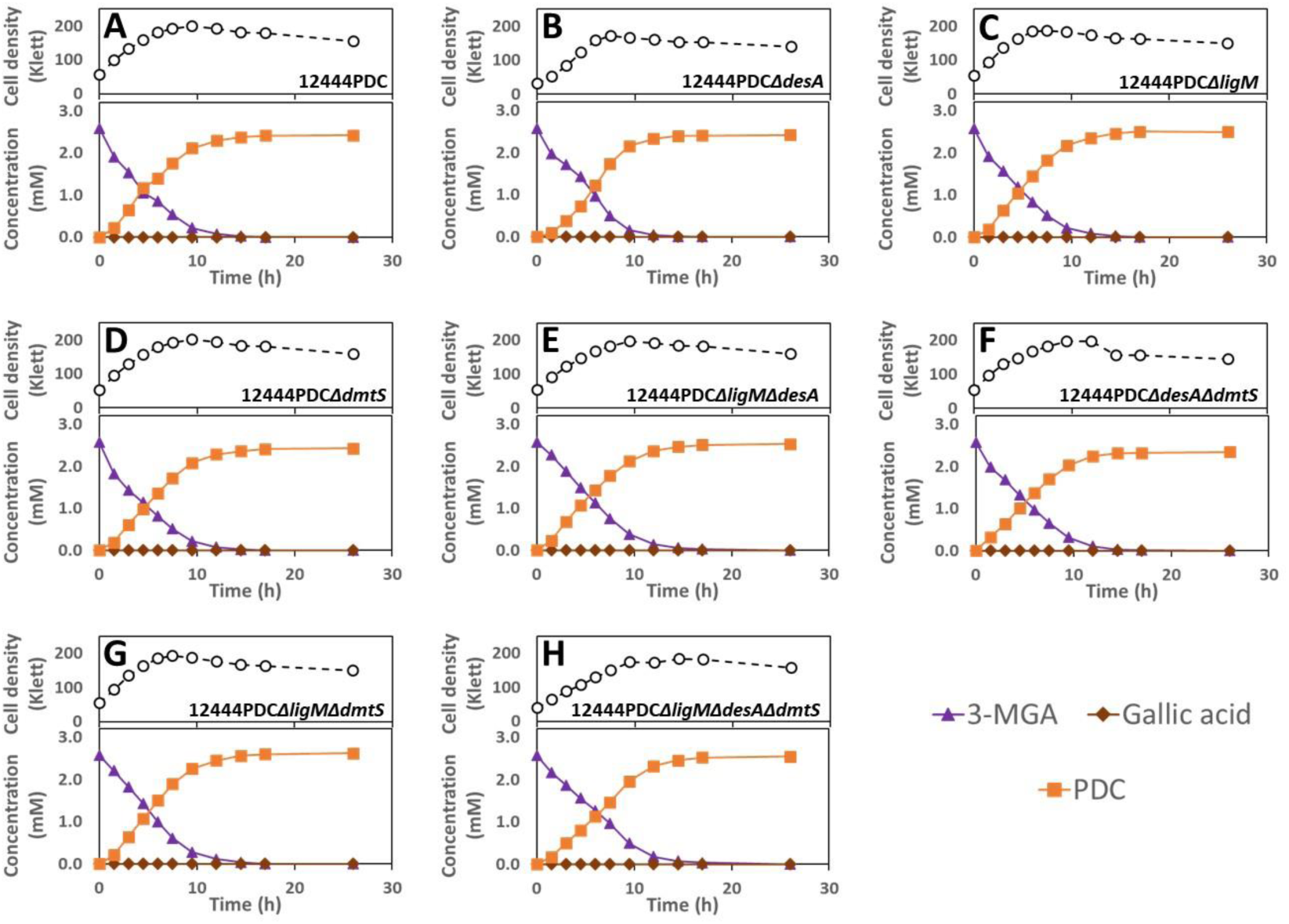
Growth and extracellular aromatic compound concentrations for the indicated *N. aromaticivorans* strains grown in minimal media supplemented with glucose and 3-MGA.

In an additional experiment with the PDC-producing strain and its derivatives, we analyzed the role of the putative *O-*demethylases in metabolism of vanillic acid. In the presence of glucose and vanillic acid, strain 12444PDC completely removed the vanillic acid from the medium and converted it into PDC with ∼96% yield (Table 2). Transient extracellular accumulation of a small amount of PCA during the course of the experiment (Figure 5A) supports the predicted role of demethylation in the degradation of vanillic acid (Figure 1). Analysis of the mutants lacking single putative *O-*demethylases (Figure 5B, 5C, 5D) revealed a decrease in the vanillic acid consumption rate when *ligM* was deleted (Figure 5C), compared to the other two strains, suggesting a role for LigM in vanillic acid metabolism. In the set of double *O-* demethylase mutants (Figure 5E, 5F, 5G), no effect was seen when *desA* and *dmtS* were deleted (Figure 5F) but a decrease in vanillic acid consumption rates was evident with all double mutants which lacked the *ligM* gene. This observation is also consistent with the reduced rate of vanillic acid degradation in the single *ligM* deletion mutant (Figure 5C). Notably, the rate of vanillic acid consumption decreased the most when *ligM* and *desA* were both deleted (Figure 5E), suggesting that DesA can partially substitute for LigM in the degradation of vanillic acid. Finally, minimal vanillic acid degradation was observed in a mutant that lacks all 3 of the putative *O-*demethylase genes (Figure 5H), consistent with the lack of LigM and DesA activity in this strain.

**Figure 5.**
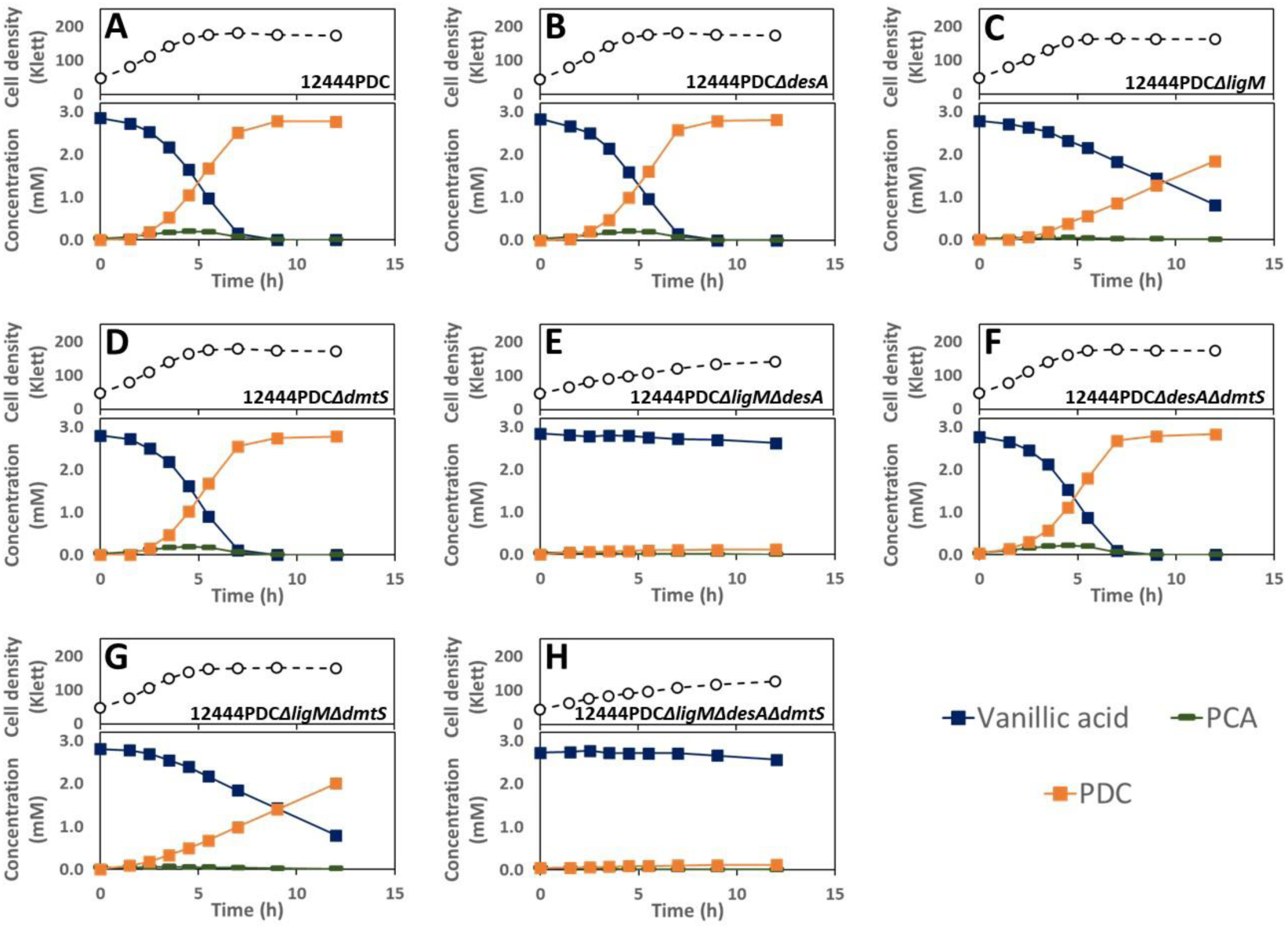
Growth and extracellular aromatic compound concentrations for the indicated *N. aromaticivorans* strains grown in minimal media supplemented with glucose and vanillic acid.

### Activity of LigM and DesA with aromatic substrates

*In vitro* experiments were performed with purified recombinant LigM and DesA proteins to test their activity with the methoxylated aromatic substrates. These assays were only performed with LigM and DesA since we have thus far been unsuccessful in purifying an active recombinant DmtS protein. The LigM and DesA homologues from *Sphingobium sp*. SYK-6 have been shown to be tetrahydrofolate (H_4_folate) dependent *O-*demethylases (25, 26), a prediction that was experimentally confirmed (Figure S1).

The recombinant LigM protein of *N. aromaticivorans* was able to convert 3-MGA into GA and vanillic acid into PCA at comparable rates under our assay conditions (Figure 6C, 6E). However, under identical conditions, the recombinant LigM protein was unable to convert a detectable amount of syringic acid into 3-MGA (Figure 6A). These results are consistent with the observations in growth experiments with mutant strains, which predicted LigM’s involvement in vanillic acid and syringic acid metabolism; in the case of syringic acid, these *in vitro* data suggest that during syringic acid metabolism, LigM does not catalyze the demethylation of syringic acid to 3-MGA, but instead demethylates 3-MGA, resulting in the small amount of syringic acid that is metabolized via GA.

**Figure 6.**
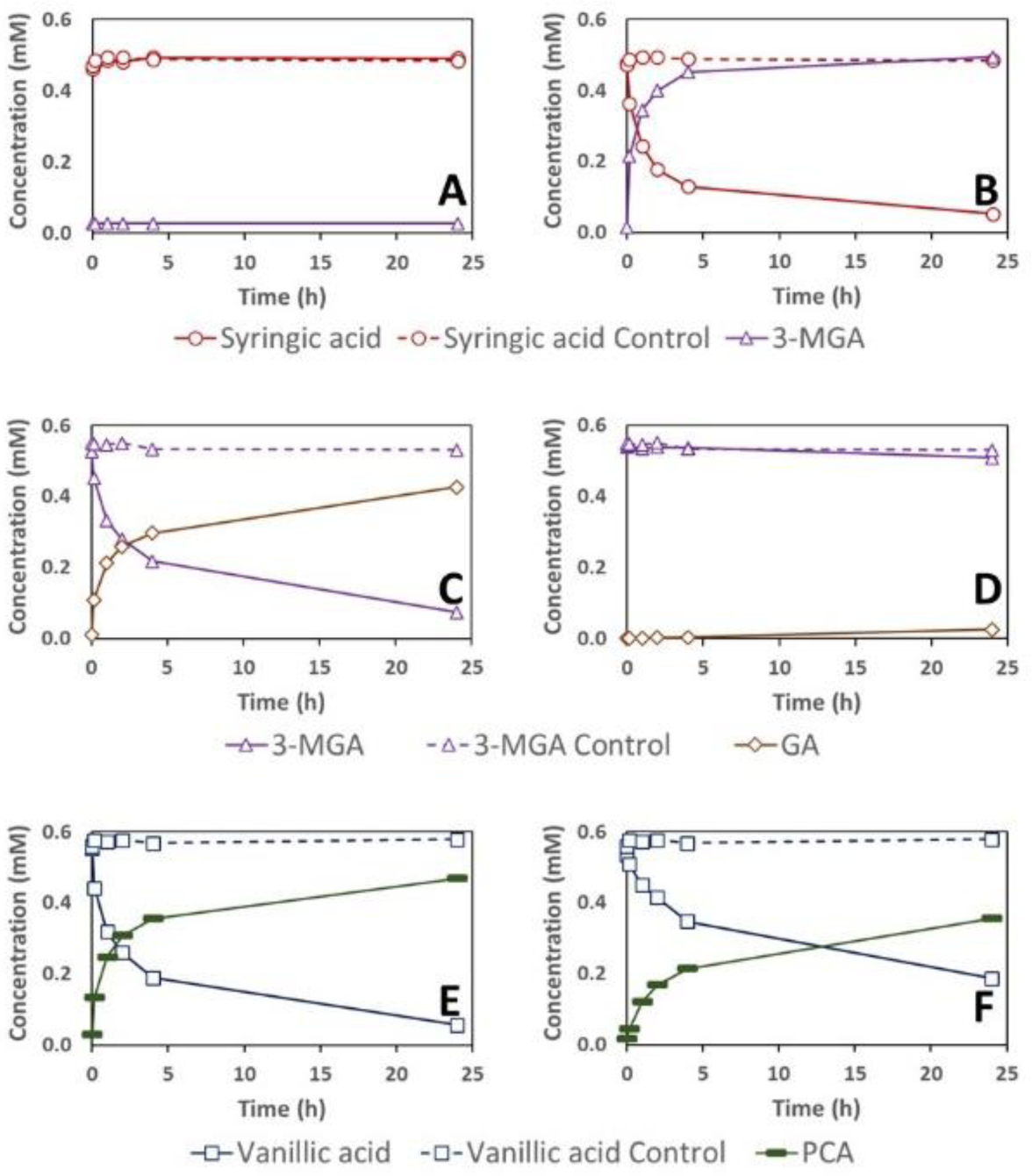
Substrate and product concentration during *in vitro* enzyme assays of LigM (Saro_2861) (panels A, C, and E) and DesA (Saro_2404) (panels B, D, and F) with syringic acid (panels A and B), 3-MGA (panels C and D), and vanillic acid (panels E and F). Concentrations in reactions lacking recombinant proteins are shown in dashed line. Reported concentrations are the average of three replicate assays. The low amounts of 3-MGA measured in the assays with LigM and syringic acid (panel A) was a trace contaminant in the syringic acid used in these assays.

We also found that the recombinant DesA of *N. aromaticivorans* demethylated both syringic acid and, somewhat more slowly, vanillic acid (Figure 6B, 6F), but was not active in demethylating 3-MGA (Figure 6D). These results support the critical role of DesA in the demethylation of syringic acid that was predicted by analyzing growth of mutant strains (Figure 2) and its potential role in vanillic acid transformation when *ligM* is deleted (Figure 5), and offer new evidence supporting the hypothesis that this enzyme does not participate in 3-MGA demethylation.

### Identification of putative aromatic ring-opening dioxygenases in *N. aromaticivorans*

The opening of the aromatic ring is an essential step in the assimilation of plant-derived phenolic compounds into intermediary metabolism (Figure 1). In *N. aromaticivorans*, the only aromatic ring-opening dioxygenase that has been previously identified is a LigAB homologue, encoded by Saro_2813 (*ligA*) and Saro_2812 (*ligB*) (15), whose subunits have ∼67% and ∼70% amino acid sequence identity with LigA and LigB of *Sphingobium* sp. SYK-6, respectively. A search of the *N. aromaticivorans* genome reveals genes that could encode another aromatic ring-opening dioxygenase encoded by Saro_1233 and Saro_1234 (hereafter referred to as *ligA2* and *ligB2*, respectively), whose subunits have amino acid sequences that are ∼33% and ∼42% identical to the LigA and LigB of *N. aromaticivorans*. We sought to test the roles of *N. aromaticivorans* LigAB and LigAB2 in the metabolism of aromatic compounds derived from plant biomass.

### Effect of deleting genes encoding putative aromatic ring-opening dioxygenases on growth of N. aromaticivorans

To evaluate the roles of LigAB and LigAB2 in the degradation of S and G phenolics by *N. aromaticivorans*, we tested growth and aromatic metabolism by mutants containing combinations of deletions in *ligAB* and *ligAB2* in the parent strain (12444*Δ1879*) and the PDC-producing strain (12444PDC) (Table 3). When cultured in the presence of either syringic acid or vanillic acid, the parent strain and strain 12444*ΔligAB2* both grew well, whereas strains with deletion of *ligAB* (12444*ΔligAB* and 12444*ΔligABΔligAB2*) were unable to grow (Figure 7). This indicates that LigAB is necessary for *N. aromaticivorans* growth on both syringic and vanillic acids, whereas LigAB2 is not.

**Table 3.** List of *N. aromaticivorans* mutant strains with deletions of putative aromatic ring cleavage dioxygenases used in this study.

| Strain | Background used for strain construction | Relevant Characteristics |
| --- | --- | --- |
| 12444ΔligAB | 12444Δ1879* | 12444Δ1879 ΔSaro_2812 ΔSaro_2813 |
| 12444ΔligAB2 | 12444Δ1879 | 12444Δ1879 ΔSaro_1233 ΔSaro_1234 |
| 12444ΔligABΔligAB2 | 12444ΔligAB | 12444Δ1879 ΔSaro_2812 ΔSaro_2813<br>ΔSaro_1233 ΔSaro_1234 |
| 12444PDCΔligAB | 12444PDC* | 12444PDC ΔSaro_2812 ΔSaro_2813 |
| 12444PDCΔligAB2 | 12444PDC | 12444PDC ΔSaro_1233 ΔSaro_1234 |
| 12444PDCΔligABΔligAB2 | 12444PDCΔligAB | 12444PDC ΔSaro_2812 ΔSaro_2813<br>ΔSaro_1233 ΔSaro_1234 |

\*Construction of the parent strain (12444 Δ1879) and the PDC-producing strain (12444PDC) is described in Perez et al. (2019) (9). See also footnote in Table 1.

**Figure 7.**
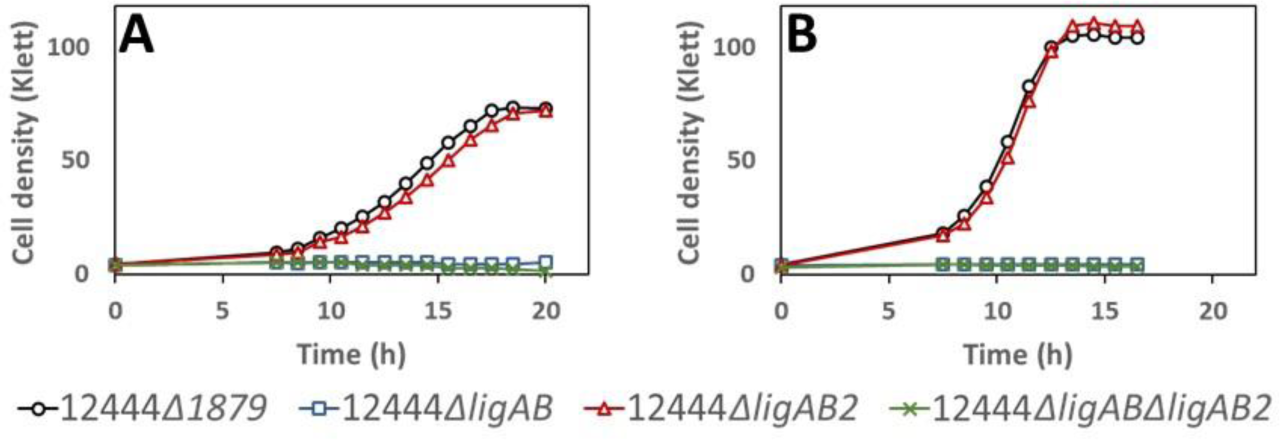
Growth of indicated *N. aromaticivorans* strains in minimal media supplemented with syringic acid (panel A) or vanillic acid (panel B).

Investigating the metabolism of syringic acid by mutants containing deletions in *ligAB* and *ligAB2* in the PDC-producing strain (12444PDC) (Figure 8), we observed that all strains consumed syringic acid, but only the mutant with intact LigAB (Figure 8B) produced PDC with a high yield (∼84%; Table 4), supporting the hypothesis that LigAB plays an important role in PDC production. The proposed pathway for PDC production from syringic acid includes ring cleavage of 3-MGA to CHMOD, which is then converted to PDC (Figure 1). The high PDC yield observed in cells in which LigAB is present (Table 4) indicates that this enzyme plays a major role in the transformation of 3-MGA. It also suggests that in *N. aromaticivorans*, 3-MGA ring opening is the primary route for 3-MGA metabolism, while demethylation of 3-MGA to GA is a secondary route responsible for only a small fraction of the metabolized 3-MGA. In support of this, the tests with the two mutants that were missing LigAB (Figure 8A, 8C) showed slower rates of syringic acid consumption and accumulation of 3-MGA and GA, the predicted metabolic intermediates in this secondary metabolic route. Strain 12444PDCΔ*ligAB* also converted only ∼2% of the syringic acid to PDC (Table 4), suggesting that LigAB2 may have some role in 3-MGA metabolism, whereas strain12444PDC*ΔligABΔligAB2* did not produce any detectable PDC (Table 4), suggesting a complete interruption of 3-MGA aromatic ring opening in the mutant that lacked both of the putative dioxygenases, LigAB and LigAB2.

**Figure 8.**
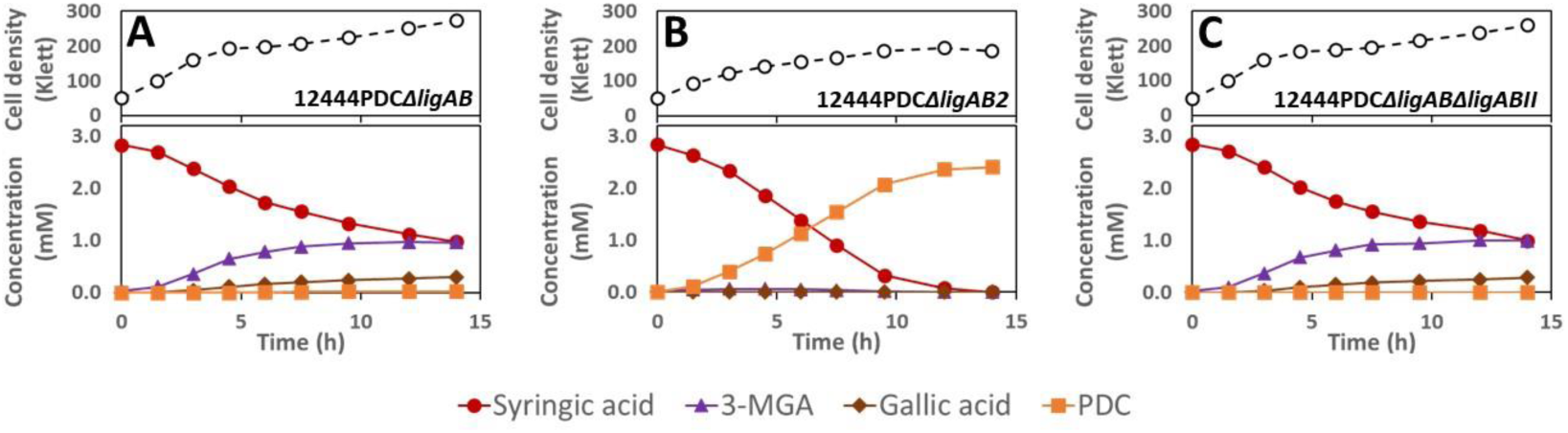
Growth and extracellular conpound concentration of indicated *N. aromaticivorans* strains cultured in minimal media supplemented with glucose and syringic acid.

**Table 4.**
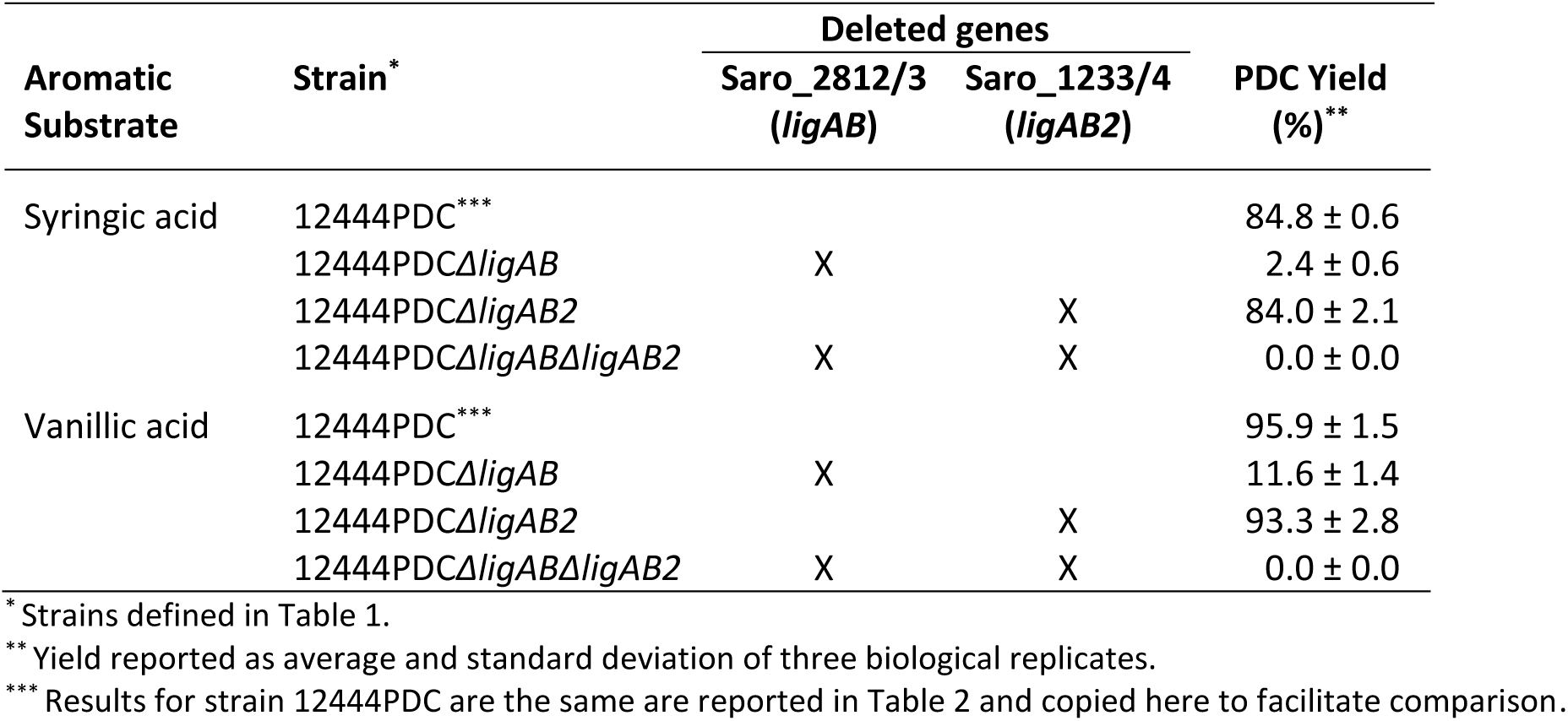
PDC yields from *N. aromaticivorans* strains in the presence of glucose plus the indicated aromatic substrate.

| Aromatic Substrate | Strain* | Deleted genes |  | PDC Yield (%)** |
| --- | --- | --- | --- | --- |
|  |  | Saro_2812/3 (*ligAB*) | Saro_1233/4 (*ligAB2*) |  |
| Syringic acid | 12444PDC*** |  |  | 84.8 ± 0.6 |
|  | 12444PDCΔ*ligAB* | X |  | 2.4 ± 0.6 |
|  | 12444PDCΔ*ligAB2* |  | X | 84.0 ± 2.1 |
|  | 12444PDCΔ*ligAB*Δ*ligAB2* | X | X | 0.0 ± 0.0 |
| Vanillic acid | 12444PDC*** |  |  | 95.9 ± 1.5 |
|  | 12444PDCΔ*ligAB* | X |  | 11.6 ± 1.4 |
|  | 12444PDCΔ*ligAB2* |  | X | 93.3 ± 2.8 |
|  | 12444PDCΔ*ligAB*Δ*ligAB2* | X | X | 0.0 ± 0.0 |

\* Strains defined in Table 1.
\*\* Yield reported as average and standard deviation of three biological replicates.
\*\*\* Results for strain 12444PDC are the same are reported in Table 2 and copied here to facilitate comparison.

Notably, in the two experiments where we observed GA accumulation (Figure 8A, 8C), the media darkened in color, which could be due to a rapid non-enzymatic transformation of GA under the conditions used for these experiments. Since the sum of the molar concentration of unreacted syringic acid plus the accumulated metabolites does not add up to 100% in either experiment, these results also suggests that the GA produced was partially degraded either abiotically or biologically.

The role of the putative ring-opening dioxygenases in vanillic acid degradation was also tested using the deletion mutants in the PDC-producing strain background (Figure 9). Both the LigAB and LigAB2 mutant derivatives of the PDC-producing strain showed degradation of vanillic acid, but only the mutant with intact *ligAB* genes produced high yields of extracellular PDC (Figure 9B, Table 4). The strains with deleted *ligAB* genes, 12444PDC*ΔligAB* (Figure 9A) and 12444PDCΔligABΔligAB2 (Figure 9C), had a slower consumption of vanillic acid, only removing ∼80% of it from the media and converting ∼48% of it into extracellular PCA over the course of the experiments. These observations suggest that LigAB has a significant role in PCA ring opening in *N. aromaticivorans*. Notably, strain 12444PDCΔligAB also converted ∼12% of the vanillic acid into PDC, whereas 12444PDCΔligABΔligAB2 did not accumulate any detectable PDC in the media, suggesting that in the absence of LigAB, LigAB2 may also function in PCA ring opening, while the absence of both LigAB and LigAB2 completely eliminates the PCA ring opening activity in *N. aromaticivorans*.

**Figure 9.**
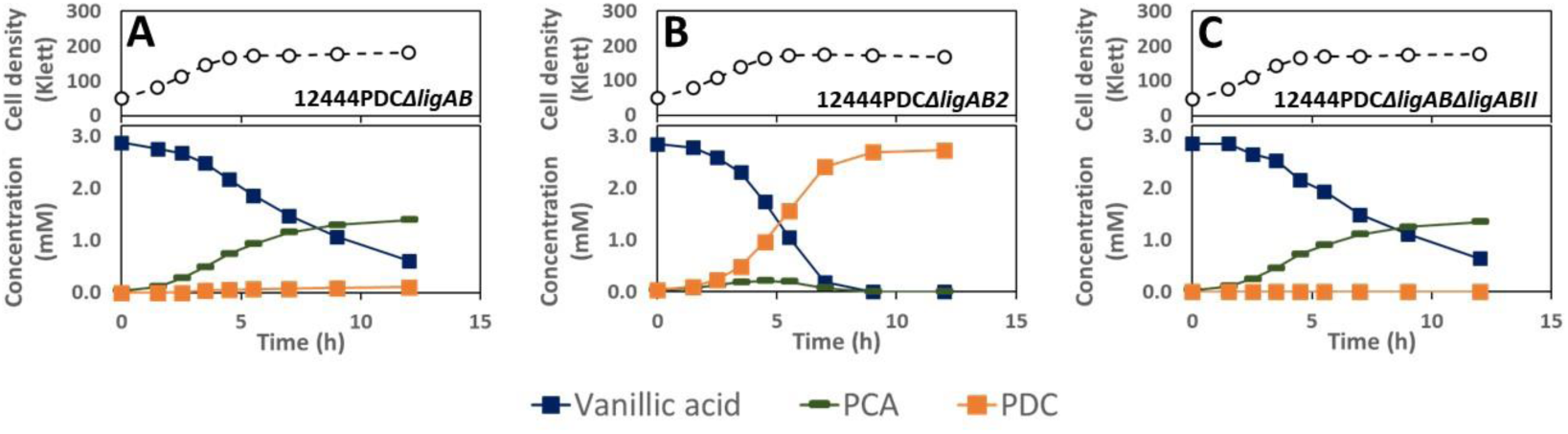
Growth and extracellular compound concentration of the indicated strains of *N. aromaticivorans* cultured in minimal media supplemented with glucose and vanillic acid.

### Activity of LigAB and LigAB2 with aromatic substrates

To investigate the predicted activities of LigAB and LigAB2 in aromatic ring opening, we purified recombinant forms of the proteins and tested them for activity *in vitro* (Figure 10). With 3-MGA as the substrate, LigAB completely removed the 3-MGA from the reaction mixture within two hours (Figure 10A), whereas LigAB2 only removed ∼20% of the 3-MGA over the course of 24 hours (Figure 10B). With each purified enzyme, two products that transiently accumulated in the assay were identified as isomers of CHMOD (See Electronic Supplementary Information), and a third product was identified as PDC. These results show that LigAB catalyzes the transformation of 3-MGA to CHMOD, and that CHMOD is subsequently transformed to PDC, as it has been proposed previously (9). They also show that LigAB2 can also convert 3-MGA to CHMOD, albeit at a slower rate than with purified LigAB under the assay conditions used, which is consistent with results from the *in vivo* analyses of defined mutants in each of the corresponding genes (Figures 7 and 8).

**Figure 10.**
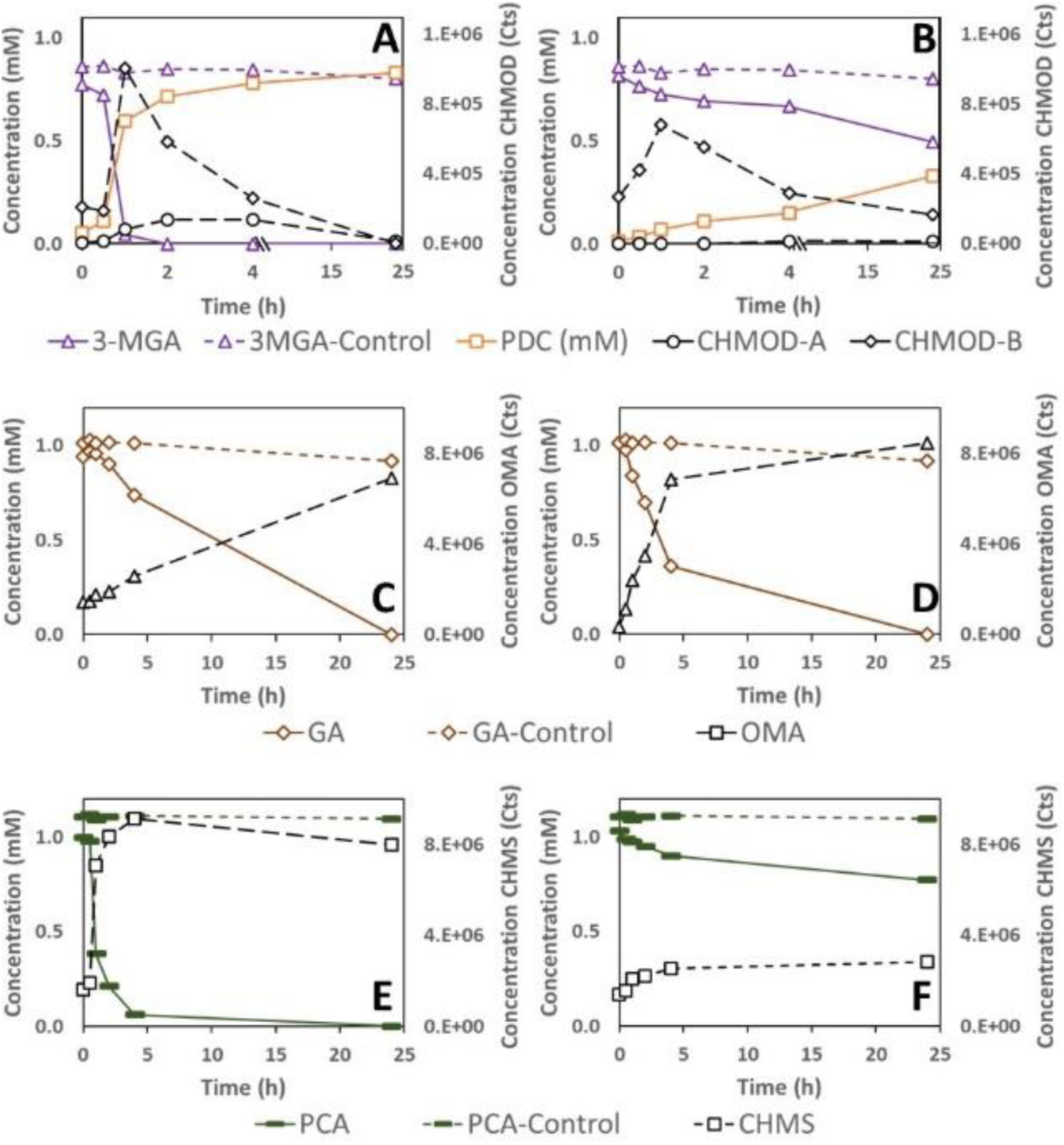
Substrate and product concentration of *in vitro* assays of LigAB (Saro_2812/3) (panels A, C, and E) and LigAB2 (Saro_1233/4) (panels B, D, and F) on 3-MGA (panels A and B), GA (panels C and D), and PCA (panels E and F). Concentration of substrates in assays performed in the absence of added recombinant protein are shown in dashed line and the same symbol and color as condition with enzyme. Samples were analyzed with HPLC-MS/MS and concentrations correspond to the average of three replicate assays.

Both LigAB and LigAB2 also showed activity when tested with GA as the substrate (Figure 10C, 10D). One compound that accumulated in these assays has been identified as OMA, the expected product of this ring opening reaction (See Electronic Supplementary Information). Under identical assay conditions, LigAB2 degraded ∼31% of the GA after 2 hours of reaction (Figure 10D), whereas LigAB degraded only ∼11% (Figure 10C). These findings suggest that of these two ring-opening dioxygenases, *N. aromaticivorans* LigAB2 has higher catalytic activity with GA than purified LigAB.

When tested with PCA as the substrate under identical assay conditions, LigAB removed ∼81% of the PCA from the reaction mixture after two hours (Figure 10E), while LigAB2 only removed ∼14% of the PCA over the same time span (Figure 10F). The reaction product from both assays was identified as CHMS (See Electronic Supplementary Information), consistent with the expected ring opening product of these enzymes with PCA as a substrate. The relative rates of PCA disappearance in assays using LigAB and LigAB2 suggests that, under the assay conditions, LigAB is more catalytically active with PCA, which is consistent with the *in vivo* experiments with the individual mutant strains (Figure 9).

## DISCUSSION

Strategies to successfully engineer efficient microbial catalysts that produce valuable compounds from chemically depolymerized lignocellulsoic biomass have several requirements, including the need for the microorganisms to funnel a heterogenous mixture of plant-derived phenolic compounds through central pathways, and the ability to genetically engineer the microorganisms to direct flow of carbon from central aromatic metabolic pathways to the production of valuable compounds. To develop these strategies, it is necessary to acquire a detailed understanding of the native aromatic metabolic pathways in the microorganisms to be used as chasses for lignin valorization. This study focuses on advancing the knowledge of native aromatic metabolism in *N. aromaticivorans*, a sphingomonad of interest because it efficiently degrades the major S, G, and H phenolic substitutents of plant biomass (9, 15) and because it is equipped with metabolic pathways for breaking down interunit linkages in lignin (17-20). The ability of *N. aromaticivorans* to efficiently degrade many aromatic compounds may be linked to having functionally redundant aromatic degradation pathways and enzymes with broad substrate specificity. While having redundant pathways and enzyme promiscuity may confer this microorganism with ecological advantages in nature, these features can create challenges or opportunites when engineering such microorganisms to produce high yields of a desired compound. For example, we reported earlier that the 12444PDC strain of *N. aromaticivorans* is able to funnel multiple lignin-derived aromatic compounds into PDC, but the PDC yields from S phenolics were lower than those from G and H phenolics (9). A possible explanation for this observation is pathway redundancy (16), which would allow *N. aromaticivorans* to channel a fraction of the S phenolics through one or more uncharacterized pathways that were not blocked in the 12444PDC strain.

Based on the published analysis of aromatic metabolism of *Sphingobium sp*. SYK-6 (Figure 1), we posited that one uncharacterized step could be the *O-*demethylation of 3-MGA to form GA with subsequent aromatic ring opening to produce OMA (Figure 1). Because *O-*demethylation and aromatic ring opening are also involved in other branches of the aromatic metabolism pathways, in this work we systematically evaluated these two functions in *N. aromaticivorans*. Below we discuss the new information derived from the integration of *in vivo* experiments with mutant strains and *in vitro* experiments with purified enzymes (summarized in Figure 11). The knowledge gained from these experiments helps us define roles for enzymes not previously described in *N. aromaticivorans* or other sphingomonads, identify functional pathway redundancy in the metabolism of S phenolics by this organism, and refine predictions of the substrate specificity of key *N. aromaticivorans* enzymes.

**Figure 11.**
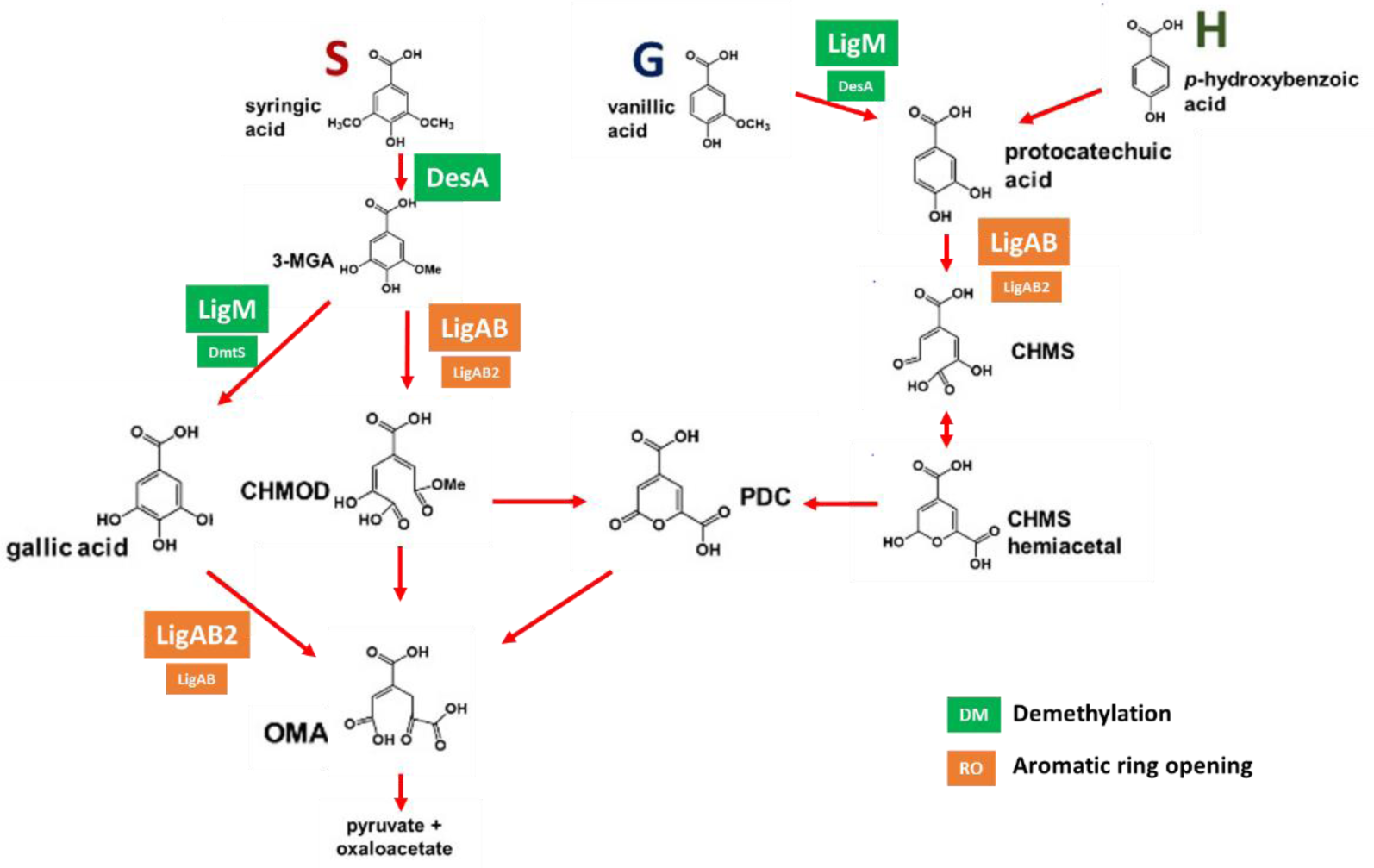
Updated pathway for metabolism of S, G, and H phenolics by *N. aromaticivorans*, with specific assignment of *O-*demethylase and aromatic ring opening dioxygenase activities based on the *in vivo* and *in vitro* experiments described in this study.

### *O-*Demethylation reactions

The two H_4_folate-dependent *O-*demethylases, DesA and LigM, and the newly identified DmtS, were each shown to have a role in the metabolism of S and G phenolics in *N. aromaticivorans* (Figure 11). While we obtained genetic evidence for the role of DmtS as an *O-*demethylase, we were not able to purify recombinant DmtS, so we lack direct information about its substrate specificity. Of known *O-* demethylases, DmtS is most similar in its predicted amino acid sequence (∼27% identical) to the monooxygenase component (subunit A) of the vanillic acid *O-*demethylase VanAB, whose function has been demonstrated in other bacteria (29, 30) and plays a key role on aromatic metabolism in *P. putida* (27, 28). In *P. putida* and other bacteria, the *vanA* gene is found in an operon with *vanB*, which encodes for the putative reductase, VanB. However, *dmtS* is not contained in a gene cluster in *N. aromaticivorans* with a gene that encodes a protein with amino acid sequence identity to a known VanB reductase. Nevertheless, we were able to use mutants lacking *dmtS* to confirm its role in aromatic metabolism in *N. aromaticivorans*.

### O-demethylation of syringic acid

Several of our results support the role of the H_4_folate-dependent *O-* demethylase DesA in demethylation of syringic acid to 3-MGA. First, the *in vitro* assays with DesA showed stoichiometric conversion of syringic acid to 3-MGA, and the *in vitro* assays with LigM preparations that were active with other aromatic compunds showed no detectable activity with this substrate (Figure 6). Second, *in vivo* experiments with the set of putative *O-*demethylase mutants showed that DesA was essential for *N. aromaticivorans* growth on syringic acid (Figure 2). Third, the experiments with the set of putative *O-*demethylase mutants constructed in the previously reported PDC-producing strain (12444PDC) showed that three out of the four mutants lacking *desA* did not degrade syringic acid (Figure 3).

To our knowledge, no enzymes other than DesA have been reported to demethylate syringic acid and produce 3-MGA in any sphingomonad. Indeed in *Sphingobium* sp. SYK-6, DesA has been proposed to be the only enzyme responsible for syringic acid *O-*demethylation and, similar to *N. aromaticivorans*, the deletion of the *desA* gene prevents the mutant strain from growing on syringic acid as the sole substrate (25). We were therefore surprised to find PDC production when *desA* was the only gene deleted in the 12444PDC background (Figure 3B). To explain this result, we hypothesize that when *N. aromaticivorans* does not use syringic acid as the growth substrate (glucose was the growth substrate with the PDC-producing strain and its derivatives), an *enzyme* other than DesA may perform the demethylation function. However, given the results from the *in vitro* assays (Figure 6), we predict that LigM is not this other *O-*demethylase, and there was no genetic evidence to link DmtS with this function.

### O-Demethylation of 3-MGA

Although *O-*demethylation of 3-MGA to GA has been described in *Sphingobium* sp. SYK-6 (26), evidence for enzymes involved in this reaction in *N. aromaticivorans* was lacking before our studies. Here, we provide several lines of evidence that 3-MGA is converted to GA in *N. aromaticivorans*, but it is not the major route of 3-MGA metabolism. Instead, our results support the hypothesis that aromatic ring opening of 3-MGA to CHMOD is the major route for the metabolism of syringic acid. First, the non-stoichiometric conversion of syringic acid to PDC in the 12444PDC strain (Figure 3A) is evidence for the existence of a secondary pathway that supports the degradation of a small fraction (∼15%) of the syringic acid in the PDC-producing strain. Second, stoichiometric conversion of syringic acid to PDC is achieved in the mutant that has an intact *desA* (necessary for the conversion of syringic acid to 3-MGA) and lacks both *ligM* and *dmtS* (Figure 3G), leading to the hypothesis that both LigM and DmtS are active in this secondary pathway. This hypothesis was further confirmed in experiments using 3-MGA as the aromatic substrate (Figure 4), where stoichiometric conversion of 3-MGA to PDC is only achieved by simultaneous deletion of *ligM* and *dmtS* (Table 2). Third, the *in vitro* assays with purified recombinant LigM (Figure 6) showed that GA was stoichiometrically produced from 3-MGA. Finally, when comparing PDC yields from the 12444PDC strain and the strains with deletion of either *dmtS* or *ligM* (Table 2), a small but reproducible increase in PDC yield occurs when *ligM* is deleted, suggesting that *in vivo* LigM is more active in 3-MGA demethylation than DmtS.

Based on a comparison of our results to that from other labs, the role of 3-MGA *O-*demethylation in the metabolism of syringic acid appears to be different in *N. aromaticivorans* and *Sphingobium sp*. SYK-6. For instance, inactivation of *ligM* in *N. aromaticivorans* did not cause a detectable effect in growth on syringic acid (Figure 2A), whereas inactivation of this gene in *Sphingobium* sp. SYK-6 has a detrimental effect on both growth rate and final cell density when cells use syringic acid (25). This suggests that between the two functionally redundant pathways for converting 3-MGA to OMA in *N. aromaticivorans*, ring opening to CHMOD carries more flow of carbon than demethylation to GA, which is the opposite of what was reported for *Sphingobium sp*. SYK-6. In addition, our results also suggest redundancy in the enzymes that demethylate 3-MGA to GA in *N. aromaticivorans*, with both LigM and DmtS having this activity. *Sphingobium sp*. SYK-6 appears to have less enzyme redundancy in this reaction, as deletion of *ligM* eliminated 3-MGA conversion to GA in cell extract experiments (25).

### O-demethylation of vanillic acid

Based on our data, we propose that LigM plays a major role in demethylation of vanillic acid, but that DesA can perform this reaction, although with reduced efficiency (Figure 11). First, the *in vitro* assays showed that both these enzymes could convert vanillic acid to PCA, with LigM having a faster degradation rate (Figure 6). Second, *in vivo* experiments with the set of *O-* demethylase mutants in the wild-type background showed that all mutants with an intact *ligM* gene grew as well as the parent strain, strains lacking *ligM* but with intact *desA* showed a detectable growth defect, and strains lacking both *ligM* and *desA* could not grow in the presence of vanillic acid (Figure 2). Consistent with these findings, our *in vivo* experiments with the set of mutants constructed in the 12444PDC background confirmed that deleting *ligM* slowed down the rates of vanillic acid degradation, deleting *desA* alone did not have an effect, but that deleting both *ligM* and *desA* prevented vanillic acid degradation (Figure 5). Finally, a role for DmtS in vanillic acid *O-*demethylation was not evident in the results of any of the experiments. These observations are consistent with the predicted role of LigM and DesA in the metabolism of vanillic acid by *Sphingobium sp*. SYK-6 (25).

### Aromatic ring opening reactions

Aromatic ring opening of phenolic compounds in sphingomonads is predicted to be catalyzed by extradiol dioxygenases (cleaving at the 4,5 position), which often exhibit broad substrate specificity (16, 31, 32). In *Sphingobium sp*. SYK-6, at least three dioxygenases are reported to be involved in metabolism of S and G compounds: LigAB, with highest activity on PCA (33), DesZ, with highest activity on 3-MGA and GA (21), and DesB, specific for GA (24). In contrast, previous analysis of *N. aromaticivorans* has implicated only one extradiol dioxygenase, LigAB, in aromatic ring opening of phenolic compounds (15). In this study, we showed that two aromatic ring opening enzymes in *N. aromaticivorans*, LigAB and LigAB2, catalyzed extradiol cleavage of 3-MGA, GA, and PCA, the three intermediates the pathways for metabolism of S, G, and H aromatics whose rings are capable of being enzymatically opened by a 4,5-dioxygense (Figure 11). In addition, our results show that LigAB has higher activity with 3-MGA and PCA than LigAB2, whereas LigAB2 has higher activity with GA than LigAB.

### Aromatic ring opening of 3-MGA

Our results support the hypothesis that LigAB is the primary enzyme for 3-MGA ring opening in *N. aromaticivorans* (15). They also provide new evidence that LigAB2 has activity with 3-MGA and could partially substitute for the role of LigAB.

The *in vitro* assays using recombinant LigAB and LigAB2 proteins with 3-MGA as a substrate showed accumulation and disappearance of the known stereoisomers of CHMOD, and accumulation of PDC (Figure 10A, 10B). *In vitro* accumulation of PDC from 3-MGA has also been observed in enzyme assays with the LigAB of *Sphingomonas sp*. SYK-6 (22). Rapid PDC accumulation in those experiments led to the proposal that PDC is a product of LigAB activity in *Sphingomonas sp*. SYK-6. Several results from our reactions of LigAB and LigAB2 from *N. aromaticivorans* with 3-MGA lead us to the different conclusion that CHMOD is the direct product of the LigAB and LigAB2 reactions with 3-MGA, and that CHMOD is non-enzymatically converted to PDC under the conditions of the assays. First, CHMOD is unstable in aqueous solution, and non-enzymatic cyclization of CHMOD to PDC at neutral pH has been reported (34). Unfortunately, the CHMOD appears to cyclize under our reaction conditions too fast for us to be able to purify this compound for determination of its molar concentration in our assays. However, the sum of the molar 3-MGA and PDC concentrations over the course of the enzymatic reactions inversely correlated with the accumulation of the CHMOD isomers (Figure S7), suggesting CHMOD is an intermediate in the conversion of 3-MGA to PDC. Third, at the end of the LigAB reaction, when 3-MGA and CHMOD are not detectable (Figure 10A), PDC accumulation reaches the initial concentration of 3-MGA in the assay, indicating the absence of any other potential products of CHMOD degradation. Finally, the results of *in vitro* assays of LigAB and LigAB2 with other substrates (Figure 10) produced products consistent with the proposed function of these enzymes as extradiol aromatic ring opening dioxygenases, and CHMOD is the expected product of such reaction when 3-MGA is the substrate (Figure 11).

### GA ring opening

The *in vitro* evidence obtained in this study indicates that LigAB2 reacts more rapidly with GA than with 3MGA or PCA, and confirms OMA as the product of GA ring opening (Figure 10). The experiments with the set of deletion mutants in the 12444PDC background (Figure 8) also provided evidence that either LigAB or LigAB2 could be responsible for metabolizing the fraction of syringic acid that is normally channeled through GA in the 12444PDC strain. However, under the conditions of our studies, the GA that accumulated in the medium apparently converted abiotically to an unknown product. Thus, further research is needed to specifically ascertain the role of LigAB and LigAB2 on GA transformation *in vivo*.

### PCA ring opening

The results of *in vitro* (Figure 10) and *in vivo* (Figure 9) experiments with vanillic acid provided evidence that the ring opening of PCA is primarily catalyzed by LigAB, but that LigAB2 could partially substitute *in vivo* when *ligAB* was deleted. Furthermore, deleting both sets of genes eliminated growth (Figure 7) and PDC production (Figure 9), indicating that no other enzyme in *N. aromaticivorans* could catalize the ring opening of PCA under our growth and media conditions.

### Implications for PDC production from lignin-derived aromatics by *N. aromaticivorans*

We have previously tested PDC production from plant-derived phenolics by *N. aromaticivorans* (9) because this native pathway intermediate is a potential building block for bio-based plastic and epoxy adhesives (35). We showed that strain 12444PDC, which contains mutations that block the conversion of PDC and CHMOD to OMA, could transform S, G, and H phenolics to PDC, but exhibited lower PDC yields from S phenolics than from G or H phenolics (9). Based on this observation we proposed that strain 12444PDC contained additional, previously unidentified, gene products that could metabolize S phenolics (9). In this study, we identified these previously unknown enzymes in the proposed alternative pathway for metabolism of syringic acid in *N. aromaticivorans* that *O-*demethylates 3-MGA to GA (Figure 11). We further showed that these same enzymes catalyze ring opening of GA to OMA (Figure 11). The activity of the enzymes in this previously unknown alternative pathway helps explain why strain 12444PDC had lower PDC yields from S phenolics, since our data suggest that ∼15% of the 3-MGA follows the GA-OMA route and does not contribute to PDC production in strain 12444PDC. Although deleting the genes in this alternative pathway could increase the yield of PDC, the broad substrate specificity of the other *N. aromaticivorans O-*demethylases described in this study implies that single gene deletions may not be sufficient to maximize PDC yields. As predicted, stoichiometric production of PDC from either syringic acid (Figure 3G) or 3-MGA (Figure 4G) was only possible with the deletion of both *ligM* and *dmtS* from the 12444PDC background. However, since *O-*demethylases are also needed in the demethylation of G phenolics, a potential side effect of *ligM* deletion (the main enzyme we found to be responsible for vanillic acid demethylation) could be the reduced conversion of G phenolics into PDC. Indeed, we observed the predicted negative effect in the rate of vanillic acid conversion when *ligM* was deleted (Figures 5C) although the PDC yield was not affected when this gene was inactivated in the 12444PDC strain (Table 2). Since DesA was shown to have activity with vanillic acid, one potential strategy to compensate for the absence of LigM could be to engineer PDC-producing strains that overexpress *desA*, as has been done with overexpression of *vanAB* in *P. putida* to accelerate vanillic acid degradation (36).

The results of our studies also make a new prediction that deleting enzymes responsible for the aromatic ring opening step does not appear to be a productive strategy for reducing the flow of substrates through this alternative *N. aromaticivorans* pathway due to the broad substrate specificity of these dioxygenases. For example, while *in vitro* assays with the newly discovered LigAB2 showed its preference for GA over 3-MGA and PCA, its inactivation did not produce measurable effects on PDC yield (Table 4). Furthermore, inactivation of both LigAB and LigAB2 would also prevent aromatic ring opening with other biomass-derived phenolic substrates. From our data, we propose that the main function of LigAB2 *in vivo* is in ring opening reactions of aromatic compounds other than S, G, and H phenolics. While *Sphingobium sp*. SYK-6 has a predicted homologue of *N. aromaticivorans* LigAB2 (encoded by SLG_37520 and SLG_37530), we are not aware of any studies that investigate its role in S, G, or H-aromatic metabolism.

In conclusion, we identified a previously unknown alternative route for syringic acid catabolism, described new enzymes that are involved in this pathway (DmtS and LigAB2), and performed a systematic genetic and enzymatic analysis of the *O-*demethylases and aromatic ring opening dioxygenases that function in *N. aromaticivorans*. The studies revealed that these *O-*demethylases and dioxygenases have broad substrate specificity suggesting they are able to participate in different reactions within individual pathways (Figure 11). The new knowledge on the metabolism of aromatic compounds by *N. aromaticivorans* obtained in this work has allowed us to design a strain (12444PDC*ΔligMΔdmtS)* with increased yield of PDC from syringic acid. It also will enable better predictions of metabolic routes that will facilitate engineering strains for improved yields of other desirable products from biomass-derived aromatics. While the interruption of 3-MGA *O-*demethylation (by deleting *ligM* and *dmtS*) in strain 12444PDC resulted in stoichiomertric PDC production from syringic acid and vanillic acid, this new strain exhibitied a reduced rate of vanillic acid degradation. This observation illustrates additional steps in aromatic acid degradation that are targets for future strain improvement.

## MATERIALS AND METHODS

### Bacterial strains, growth media and culturing conditions

Two variants of *N. aromaticivorans* DSM12444, strains 12444*Δ1879*, which lacks the gene Saro_1879 (putative *sacB*; Saro_RS09410) (17), and strain 12444*ΔligIΔdesCD* (also called 12444PDC), which lacks the genes Saro_1879, Saro_2819 (Saro_RS14300), Saro_2864 (Saro_RS14525), and Saro_2865 (Saro_RS14530) (9) were used as parent strains to generate the mutant strains shown in Table 1.

All genetic modifications were carried out using a variant of the plasmid pK18*mobsacB* (38), which contains *sacB* and a kanamycin resistance gene. The gene deletions were performed as previously described (9), with the details of the processes used here described in the Electronic Supplementary Information. All primers, plasmids, and *Escherichia coli* strains used for cloning and protein expression are listed in supplemental Tables S1, S2, and S3, respectively.

*E. coli* cultures were grown in LB media containing 50 µg ml^-1^ kanamycin at 37 °C. *N. aromaticivorans* cultures were grown in SMB media (see Supporting Information of Kontur et al. (2018) (17) for recipe) supplemented with the indicated carbon source at 30 °C. For routine culture and storage, SMB was supplemented with 1 g L^-1^ glucose. For constructing mutants, SMB was supplemented with 1 g L^-1^ glucose and either 50 µg ml^-1^ kanamycin, or 100 g L^-1^ sucrose as necessary. Cell density was monitored using a Klett-Summerson photoelectric colorimeter with a red filter.

### *N. aromaticivorans* growth experiments

*N. aromaticivorans* cultures were grown overnight in SMB medium supplemented with 1 g L^-1^ glucose. Cultures were diluted 1:1 with fresh medium containing 1 g L^-1^ glucose and incubated for 1 hour to resume cell growth. Cells from 2 ml of the growing culture were pelleted (5 min at 2,300 × g), washed with 1 ml SMB media without carbon, then resuspended into 600 µl SMB media with no added carbon. Growth experiments were initiated by adding 80 µl of the cell suspension into 8 ml fresh SMB media supplemented with either 3 mM syringic acid or 3 mM vanillic acid. Cultures were grown aerobically in 20 ml test tubes, shaken at 200 rpm at 30°C. Each experiment was repeated 3 times.

### *N. aromaticivorans* extracellular metabolite analysis

Bacterial cell cultures were grown overnight in 20 ml SMB media supplemented with 1 g L^-1^ glucose, then reactivated by adding 2 ml fresh media containing 1 g L^-1^ glucose and incubated for 1 hour. Experiments were initiated by inoculating 5 ml active culture into 15 ml fresh SMB media supplemented with 5 mM glucose and either 4 mM syringic acid, 4 mM 3-MGA, or 4 mM vanillic acid. Cultures were grown aerobically in 125 mL conical growth flasks, shaken at 200 rpm at 30°C. Samples were collected periodically, filtered (through 0.22 µm pores) to remove cells, and immediately analyzed by HPLC-MS to monitor the extracellular aromatic compounds. Each experiment was repeated 3 times.

### Recombinant enzyme expression and purification

Genes Saro_2812/3 (*ligAB*), Saro_1233/4 (*ligAB2*), Saro_2861 (*ligM*), and Saro_2402 (*desA*) from *N. aromaticivorans* were independently cloned into the plasmid pVP302K (19), which incorporates a His_8_-tag to the N-terminus of the translated transcripts connected via a tobacco etch virus (TEV) protease recognition site (see Electronic Supplementary Information for plasmid construction details). The expression plasmids were transformed into *E. coli* B834 (39, 40) containing plasmid pRARE2 (Novagen, Madison, WI) and transformants grown in ZYM-5052 Autoinduction Medium (41) containing 50 µg ml^-1^ kanamycin and 20 µg ml^-1^ chloramphenicol. Recombinant proteins were purified by passing crude *E. coli* lysates through a nickel-nitrilotriacetic acid (Ni^2+^-NTA) column as described previously (17). His_8_ tags were cleaved from recombinant proteins using TEV protease, and proteins were passed again though a Ni-NTA column to remove the cleaved His_8_ tag and the TEV protease (which contains its own His tag).

### *In vitro* aromatic ring-opening dioxygenase assays

Preliminary experiments with recombinant LigAB (Saro_2812/3) and LigAB2 (Saro_1233/4) purified in the presence of air suggested that the enzymes were catalytically inactive, as reported for other homologues of these proteins (42). We thus separately mixed LigAB and LigAB2 with Reactivation Buffer (Buffer A, containing 20 mM Na_2_HPO_4_, 20 mM KH_2_PO_4_, 1 mM Fe_2_SO_4_ and 1 mM ascorbic acid, and prepared anaerobically) to a concentration of 2 μM enzyme active sites under anaerobic conditions, and incubated them anaerobically at 30°C for 21 hours to reactivate the enzymes. Three solutions containing Buffer B (20 mM Na_2_HPO_4_, 20 mM KH_2_PO_4_, and 1 mM ascorbic acid) and either 2 mM 3-MGA, 2 mM PCA or 2 mM GA were prepared in the presence of air. Enzyme assays were performed in 2ml polypropylene vials inside an anaerobic chamber at 30°C by mixing 750 μL of reactivated enzyme in Buffer A with 750 μL of aromatic substrate in Buffer B. As a control, we mixed 750 μL of Buffer A without enzyme with 750 μL each aromatic substrate in Buffer B. After 30 min, the vials were exposed to the ambient atmosphere outside of the anaerobic chamber for 10 min to expose the reactions to the O_2_ predicted to be a substrate for the ring-opening reaction, then transferred back into the anaerobic chamber. Exposure to O_2_ was repeated at 2 hours and 4 hours after the assay was initiated. Samples (250 μL) were collected at time zero, at 30 minutes and at 1, 2, 4, and 24 hours, and immediately mixed with 50 μL of 1N HCl to terminate the reaction, then analyzed with HPLC-MS to quantify substrate disappearance and formation of any products. Identification of OMA, CHMOD, and PDC were performed by GC-MS of samples extracted with ethyl acetate immediately after collection. CHMS was converted into 2,4-pyridinedicarboxylic acid (PDCA) by the addition of (NH_4_)_2_SO_4_ (33), which was then identified by HPLC/UV. Assays were performed in triplicate.

### Aromatic *O-*demethylase enzyme assays

Enzyme assays were performed in triplicate under anaerobic conditions at 30°C (because of the expected O_2_ sensitivity of H_4_folate). Recombinant LigM (Saro_2861) and DesA (Saro_2404) were separately mixed with Buffer C (20mM Na_2_HPO_4_, 20mM KH_2_PO_4_, and 1 mM or 2 mM H_4_folate, and prepared anaerobically) to concentrations of 2 μM enzyme active sites under anaerobic conditions. Three independent solutions containing Buffer D (20mM Na_2_HPO_4_ and 20mM KH_2_PO_4_) and either 1 mM 3-MGA, 1 mM PCA or 1 mM GA were prepared aerobically. Enzyme reactions were initiated in 2 ml polypropylene vials inside an anaerobic chamber at 30°C by mixing 750 μL of an enzyme in Buffer C with 750 μL an aromatic substrate in Buffer D. A control was run by mixing 750 μL of Buffer C without enzyme with 750 μL each aromatic substrate in Buffer D. Samples (250 μL) were collected at time zero, at 30 minutes and at 1, 2, 4, and 24 hours, and immediately mixed with 50 μL of 1N HCl to stop the reaction, then analyzed with HPLC-MS to quantify substrate disappearance and product formation.

### Analysis of extracellular metabolites and enzyme reaction products

All culture supernatants and *in vitro* enzyme assay samples were filtered (0.2 µm) prior to chemical analysis. Quantitative analyses of aromatic compounds were performed on a Shimadzu triple quadrupole liquid chromatography mass spectrometer (LC-MS) (Nexera XR HPLC-8045 MS/MS). Reverse-phase HPLC was performed using a binary gradient mobile phase consisting of Solvent A (0.2% formic acid in water) and methanol (gradient profile shown in Figure S7), and a Kinetex F5 column (2.6 μm pore size, 2.1 mm ID, 150 mm length, P/N: 00F-4723-AN). All compounds were detected by multiple-reaction-monitoring (MRM) and quantified using the strongest MRM transition (Table S6).

Identification of OMA and PDC was performed by gas chromatography-mass spectrometry (GC-MS). Sample aliquots (150 μL) were acidified with HCl to pH < 2, and ethyl acetate extracted (3 × 500 μL). The three extraction samples were combined, dried under a stream of N_2_ at 40 °C, and derivatized by the addition of 150 μL of pyridine and 150 μL of N,O-bis(trimethylsilyl)trifluoro-acetamide with trimethylchlorosilane (99 : 1, w/w, Sigma) and incubated at 70 °C for 45 min. The derivatized samples were analyzed on an Agilent GC-MS (GC model 7890A, MS Model 5975C) equipped with a (5% phenyl)-methylpolysiloxane capillary column (Agilent model HP-5MS). The injection port temperature was held at 280 °C and the oven temperature program was held at 80 °C for 1 min, then ramped at 10 °C min^−1^ to 220 °C, held for 2 min, ramped at 20 °C min^−1^ to 310 °C, and held for 6 min. The MS used an electron impact (EI) ion source (70 eV) and a single quadrupole mass selection scanning at 2.5 Hz, from 50 to 650 m/z. The data was analyzed with Agilent MassHunter software suite.

The product of PCA aromatic ring opening, predicted to be CHMS, was analysed by its conversion into 2,4-pyridinedicarboxylic acid (PDCA). 100 μl of sample were pH neutralized by the addition of 10 μl NaOH 1.67 N solution. In addition, 5 μl (NH_4_)_2_SO_4_ 10% solution was added and then incubated at room temperature for 24 h. Samples were analyzed by HPLC-UV using the same HPLC conditions described above. Eluent was analyzed for light absorbance between 190 and 400 nm by a Shimadzu SPD-M20A spectrophotometer.

### Chemicals

All SMB media reagents, gallic acid, and vanillic acid were purchased from SigmaAldrich (St Louis, MO). Syringic acid was purchased from TCI (Tokyo Chemical Industry)-America (Portland, OR). 3-MGA was purchased from Carbosynth (Berkshire, UK). Protocatechuic acid was purchased from Sigma Aldrich (St Louis, MO).

## ACKNOWLEDGENTS

This work was supported by U.S. Department of Energy (DOE) Great Lakes Bioenergy Research Center grant (DOE Office of Science BER DE-SC0018409). Additional funding from the Chilean National Commission for Scientific and Technological Research (CONICYT) as a fellowship to Jose M. Perez is also acknowledged.

